# Basement membrane extract potentiates the endochondral ossification phenotype of bone marrow-derived mesenchymal stem cell-based cartilage organoids

**DOI:** 10.1101/2023.12.11.571194

**Authors:** Hinako Notoh, Satoshi Yamasaki, Nobuaki Suzuki, Atsuo Suzuki, Shuichi Okamoto, Takeshi Kanematsu, Naruko Suzuki, Akira Katsumi, Tetsuhito Kojima, Tadashi Matsushita, Shogo Tamura

## Abstract

Endochondral ossification is a developmental process in the skeletal system and bone marrow of vertebrates. During endochondral ossification, primitive cartilaginous anlages derived from mesenchymal stem cells (MSCs) undergo vascular invasion and ossification. *In vitro* regeneration of endochondral ossification is beneficial for research on the skeletal system and bone marrow development as well as their clinical aspects. However, to achieve the regeneration of endochondral ossification, a stem cell-based artificial cartilage (cartilage organoid, Cart-Org) that possesses an endochondral ossification phenotype is required. Here, we modified a conventional 3D culture method to create stem cell-based Cart-Org by mixing it with a basement membrane extract (BME) and further characterized its chondrogenic and ossification properties. BME enlarged and matured the bone marrow MSC-based Cart-Orgs without any shape abnormalities. Histological analysis using Alcian blue staining showed that the production of cartilaginous extracellular matrices was enhanced in Cart-Org treated with BME. Transcriptome analysis using RNA sequencing revealed that BME altered the gene expression pattern of Cart-Org to a dominant chondrogenic state. BME triggered the activation of the SMAD pathway and inhibition of the NK-κB pathway, which resulted in the upregulation of *SOX9*, *COL2A1*, and *ACAN* in Cart-Org. BME also facilitated the upregulation of genes associated with hypertrophic chondrocytes (*IHH*, *PTH1R,* and *COL10A1*) and ossification (*SP7*, *ALPL*, and *MMP13*). Our findings indicate that BME promotes cartilaginous maturation and further ossification of bone marrow MSC-based Cart-Org, suggesting that Cart-Org treated with BME possesses the phenotype of endochondral ossification.

**Highlights:**

- Basement membrane extract (BME) enlarges MSC-based Cart-Org.
- BME activates the SMAD pathway and inhibits the NK-kB pathway of the Cart-Org.
- BME promotes cartilaginous maturation and further ossification of Cart-Org.

## Introduction

Cartilage is a connective tissue composed of chondrocytes and several extracellular matrices (ECMs), categorized into three types based on its compositions: hyaline cartilage at the articular surface of joints and in the nasal septum; elastic cartilage in the pinna, larynx, and epiglottis; and fibrocartilage in the intervertebral discs, sacroiliac joints, pubic symphysis, and costochondral joints (1). Chondrogenesis is a cellular process in which mesenchymal stem cells (MSCs) differentiate into chondrogenic progenitors and their derivatives to form cartilage (2). Chondrogenic progenitors enter a series of differentiation/maturation steps following proliferating chondrocytes, pre-hypertrophic chondrocytes, and hypertrophic chondrocytes. Chondrocytes secrete cartilaginous ECMs components, such as collagen type II Alpha1 (COL2A1) and proteoglycans [for example, aggrecan (ACAN)] in proliferating chondrocytes and collagen type X in pre-phypertrophic/hypertrophic chondrocytes. Chondrocytes enter a late hypertrophic state, producing a mineralized environment with calcium salts (3).

Cartilage is crucial for bone and marrow development. Long bones and the vertebrate skeleton are formed by a multistep process called endochondral ossification (4). During endochondral ossification, MSCs condense to form a dense mass and undergo chondrogenesis, generating a primitive avascular cartilaginous anlage. Vascular invasion into the cartilaginous anlage triggers the formation of an ossification center (5). In the process of ossification center formation, cartilage-specific ECMs are progressively degraded, generating a vascularized marrow space (3). In the long bone, there are different temporal-spatial mechanisms of endochondral ossification: an embryonal primary ossification center (POC) in the diaphysis and a postnatal secondary ossification center (SOC) in the epiphysis (4,6–8).

Regeneration of endochondral ossification is a useful experimental tool for research on skeletal system and bone marrow development. However, to regenerate the endochondral ossification *in vitro*, it is necessary to create a fully differentiated/matured cartilage template via stem cell chondrogenesis. To obtain an appropriate stem cell-based cartilage template for endochondral ossification, we modified a conventional three-dimensional (3D) culture method to create stem cell-based artificial cartilage by mixing basement membrane extract (BME) and characterized its chondrogenic properties. The results of this study provide fundamental knowledge for the establishment of an artificial cartilage for the regeneration of endochondral ossification *in vitro*.

## Results

### BME-mixed 3D pellet culture enlarges human bone marrow (BM) MSC-based cartilage-organoid (Cart-Org)

Three-dimensional pellet culture is a general system for the creation of stem cell-based artificial cartilage, i.e., cartilage-organoid (Car-Org) (9,10). In this study, human bone marrow-derived mesenchymal stem cells (BM-MSCs) were used as a cellular source of Cart-Org because human BM-MSCs possess potent properties that occur in endochondral ossification (11). In the conventional 3D pellet culture method, BM-MSCs are directly suspended in a chondrogenic medium and condensed by centrifugation, followed by the vertical induction of chondrogenesis (Fig. 1A). As a modified method, we arranged the pellet preparation sequence and obtained a gelated BM-MSC/BME mixture. After 21 days of cultivation, Cart-Orgs mixed with BME were constantly enlarged (Fig. 1B and C). The sectional area and diameter of Cart-Org with BME were considerably higher than those of Cart-Org without BME (Fig. 1D and E). Their circularity and roundness did not differ with or without BME (Fig. 1F and G). These observations indicate that the BME grows to the size of a Cart-Org without a shape anomaly.

**Figure 1.**
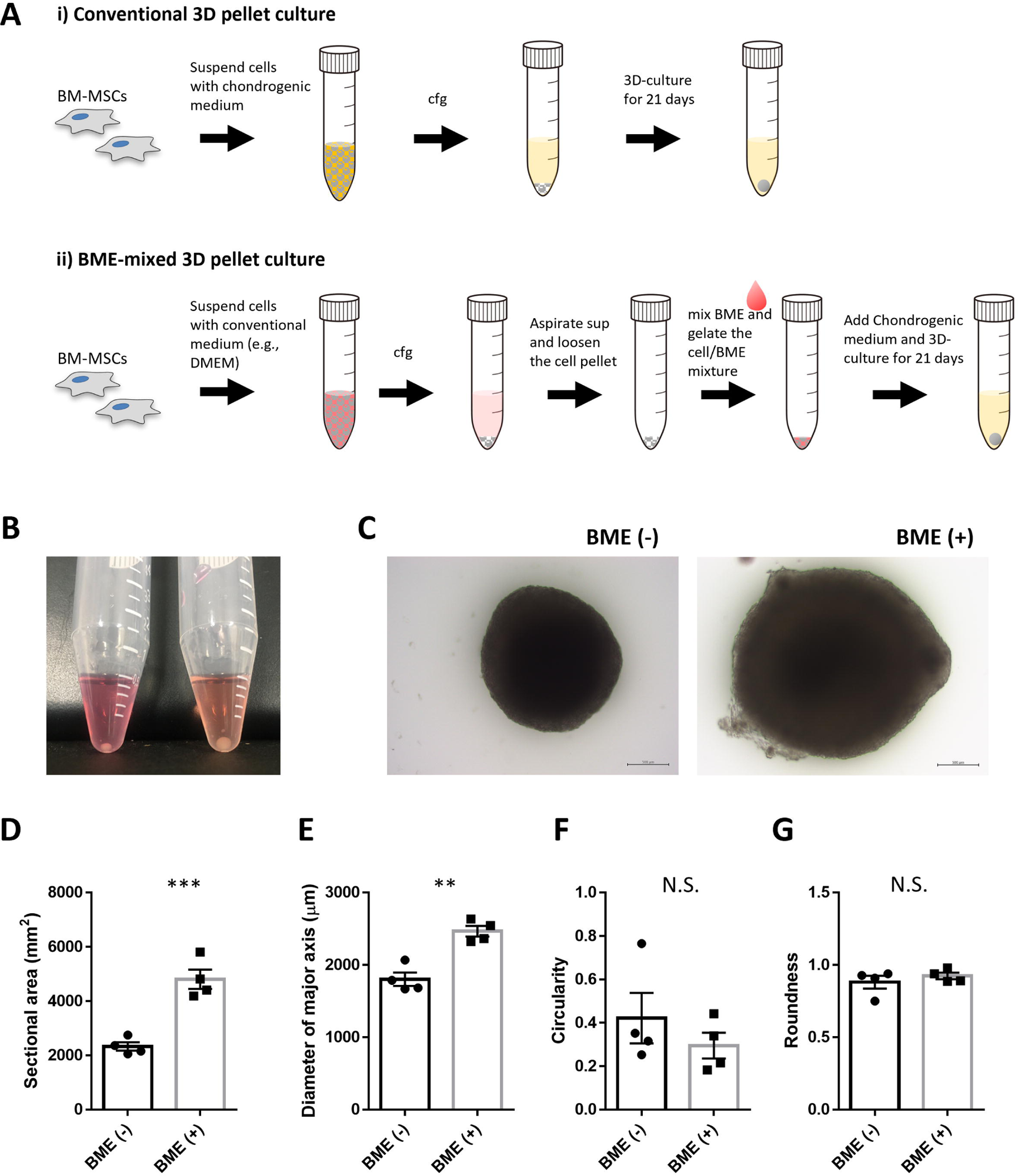
BME-mixed 3D pellet culture enlarges human BM MSC-based cartilage-organoid (Cart-Otg) without shape abnormality. A, Graphical scheme of a conventional 3D pellet culture (i) and BME-mixed 3D pellet culture (ii). B, representative appearances of human BM MSC-based Cart-Org without BME (left) and with BME (right). C, representative microscopic images of the human BM-MSC-based Cart-Org with or without BME. The scale bar indicates 500 μm. D-G, Aspect parameters of the Cart-Org with or without BME. The parameters used to evaluate aspect of the Cart-Org, including the sectional area (D), the diameter of the major axis (E), the circularity (F), and the roundness (G), were measured using Image J. **p < 0.01. ***p < 0.001. N.S. indicates a non-significant difference. Statistical analysis was performed via unpaired t test (n = 4 per group). The error bars represent SEMs. BME, basement membrane extract; BM MSC, bone marrow-derived mesenchymal stem cell; cfg, centrifugation; SEM, standard error of the mean.

### BME promotes cartilage development of BM MSC-based Cart-Org

To validate the chondrogenic quality of BM MSC-based Cart-Org with BME, we investigated its histological features by staining with hematoxylin and eosin (HE) and Alcian blue (Fig. 2). Alcian blue is a specific stain for the acidic mucopolysaccharides present in hyaline cartilage. In culture conditions without BME, Cart-Orgs showed cartilaginous phenotypes (chondrocyte hypertrophy and positive staining with Alcian blue) on day 21, but not on days 7 and 14. In the BME-mixed condition, interstitial BME depositions still existed in the cell condensate on day 7. Cartilaginous phenotypes were clearly observed in Cart-Org on day 14 and were further accentuated on day 21. These observations indicate that BME facilitates chondrogenic differentiation and maturation in MSC-based Cart-Orgs.

**Figure 2.**
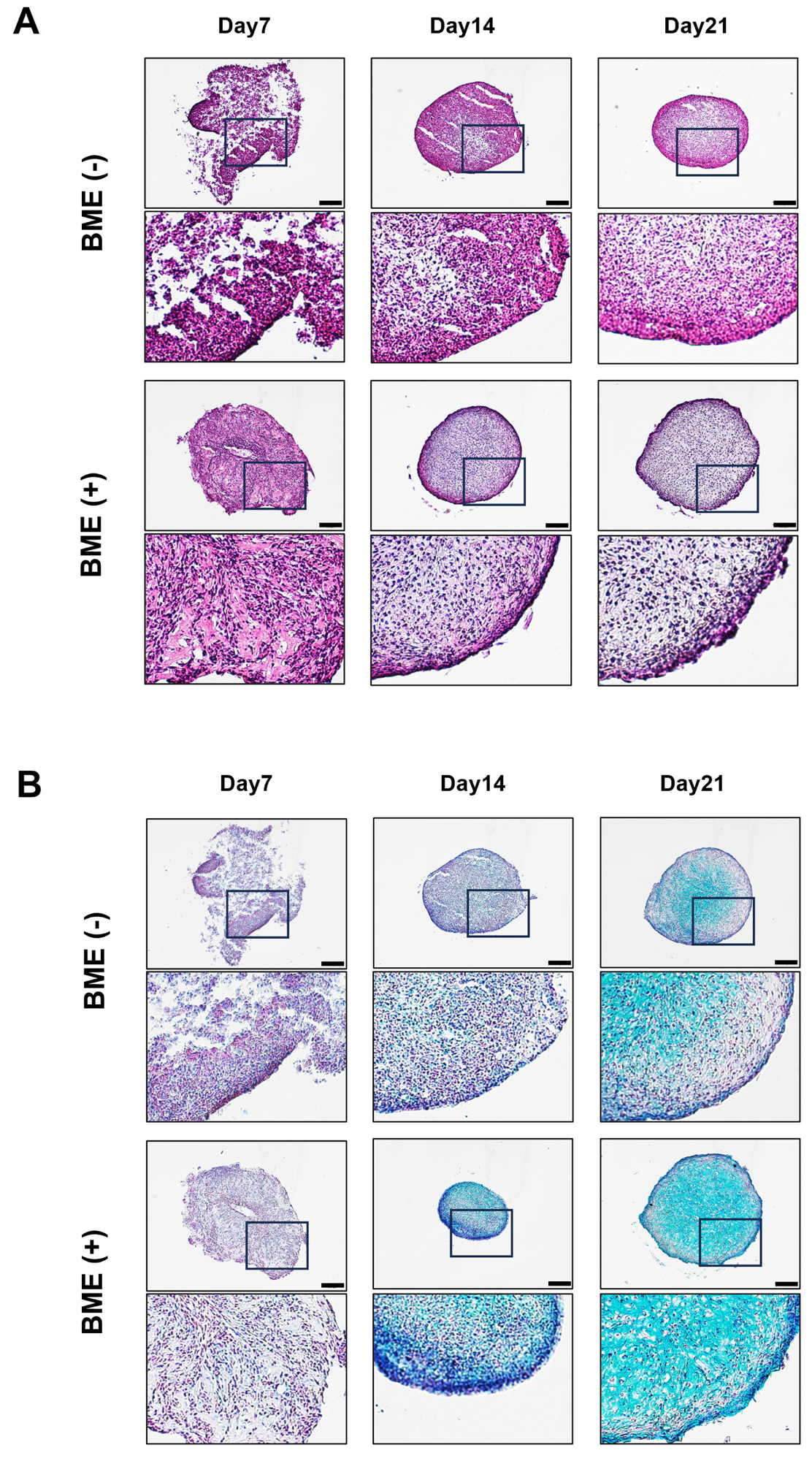
Histological analysis of the BM MSC-based Cart-Org with or without BME. A and B, representative images of the BM MSC-based Cart-Orgs stained with HE (A) and Alcian blue (B). The scale bar indicates 200 μm. BME, basement membrane extract; BM MSC, bone marrow-derived mesenchymal stem cell; Cart-Org, cartilage-organoid.

To analyze progressive chondrogenesis in the BME-mixed condition, we comprehensively investigated the gene expression of Cart-Org with BME using RNA sequencing (RNA-seq). The BME-mixed 3D culture system altered the transcriptomic pattern of the Cart-Org (Fig. 3A). On day 7, 194 upregulated and 89 downregulated genes were detected in the Cart-Org with BME (Fig. S1A, criteria: p < 0.05, fold change > 2 or < -2). The number of upregulated or downregulated genes increased throughout the course of cultivation (219 upregulated and 102 downregulated genes on day 14 and 431 upregulated and 365 downregulated-genes on day 21, Fig. S1B and C, criteria: p < 0.05, fold change > 2 or < -2). Principle component analysis (PCA) denoted the differentiation trajectory of each Cart-Org series over time (Fig. 3B). In this context, chondrogenic development appears to describe the right-to-left trace of the PC1 axis. The Cart-Org cluster without BME on day 21 (Cluster 3 in Fig. 3B) was found at approximately the same point as that of Cart-Org with BME on day 14 (Cluster 5 in Fig. 3B). This result suggests that the developmental status of Cart-Org with BME on day 14 was identical to that of Cart-Org without BME on day 21.

**Figure 3.**
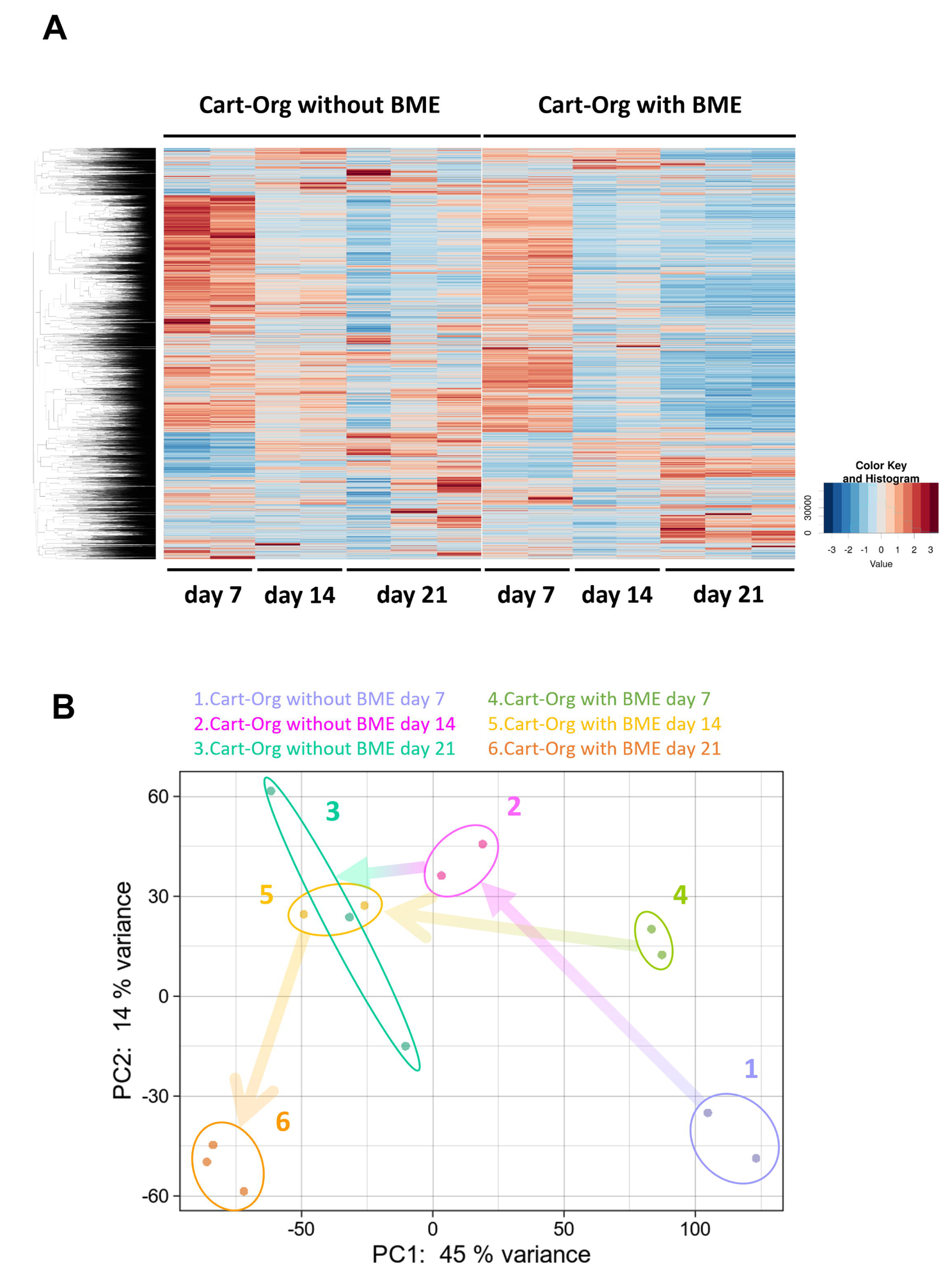
Comprehensive transcriptome analysis of the BM MSC-based Cart-Org with and without BME. A and B, each type of BM MSC-based Cart-Org on cultivation days 7, 14, and 21 was subjected to RNA-seq and their transcriptomic data of differentially expressed genes was analyzed by heat map analysis (A) and principal component analysis (B). BME, basement membrane extract; BM MSC, bone marrow-derived mesenchymal stem cell; Cart-Org, cartilage-organoid.

### BME induces SMAD pathway activation and NF-κB inhibition, promoting chondrogenesis and further ossification of the BM MSC-based Cart-Org

To investigate the mechanism by which the BME-mixed 3D pellet culture system promotes chondrogenic differentiation in the BM MSC-based Cart-Org, we performed a transcriptome-based signaling pathway analysis using a dataset of Cart-Org with/without BME on day 7. Ingenuity pathway analysis (IPA) upstream regulator analysis uncovered the top 10 activated and inhibited upstream regulators in the Cart-Org with BME on day 7 (Fig. 4A, significance criteria: p < 0.05, Z-score > 2 or < -2). Bone morphogenic protein 2 (BMP2), SRY-box transcription factor 9 (SOX9), BMP4, cytokine-inducible SH2-containing protein (CISH), N-acylsphingosine amidohydrolase 1 (ASAH1), interleukin 10 receptor subunit alpha (IL10RA), Sma and Mad related protein family 1 (SMAD1), RB transcriptional corepressor 1 (RB1), bone morphogenetic protein receptor type 1A (BMPR1A), and sprouty RTK signaling antagonist 2 (SPRY2) were predicted as the activated regulators in the Cart-Org with BME. In contrast, epidermal growth factor (EGF), interleukin 1b (IL1B), hepatocyte growth factor (HGF), tumor necrosis factor (TNF), nuclear protein 1 (NUPR1), Kirsten rat sarcoma viral oncogene homolog (KRAS), progesterone receptor (PGR), triggering receptor expressed on myeloid cells 1 (TREM1), platelet derived growth factor subunit B/B (PDGF BB), and extracellular signal-regulated kinase 1/2 (ERK1/2) were predicted as the inhibited regulators in the Cart-Org with BME. Each pathway signaling toward *SOX9, COL2A1*, and *ACAN* are shown in Fig. S2 and S3.

**Figure 4.**
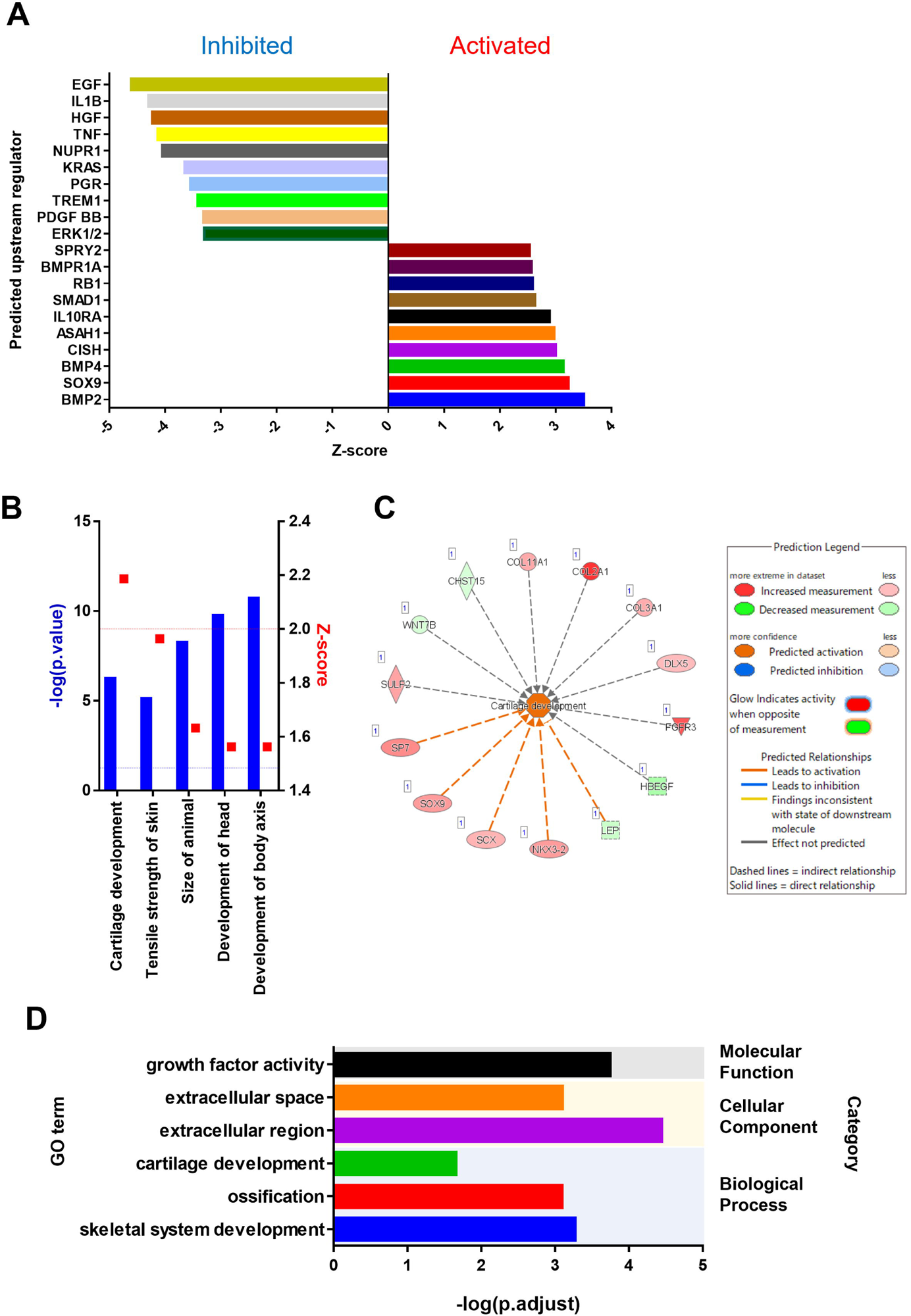
Ingenuity pathway analysis (IPA) and gene ontology (GO) analysis indicate that BME simulation promotes cartilage development and further ossification of the BM MSC-based Cart-Org. A–C, IP upstream regulator analysis using a transcriptomic dataset of the BM MSC-based Cart-Org with/without BME on day 7. A, the chart of the significant top 10 activated and inhibited upstream regulators in the BM MSC-based Cart-Org with BME. The significance of the data was determined via the following criteria: p < 0.05, Z-score > 2, or < -2. B, the chart of the IP downstream effects analysis using a dataset of the BM MSC-based Cart-Org on day 7. The annotated biological activities in the BM MSC-based Cart-Org with BME are shown. The significance of the data was determined via the following criteria: p < 0.05 and Z-score > 2. C, a molecular network related to cartilage development in the BM MSC-based Cart-Org with BME. D, GO analysis using a transcriptomic dataset of the BM MSC-based Cart-Org with/without BME on day 21. The enriched gene sets (GO terms) of the biological process, cellular component, and molecular function are shown. The significance of the data was determined via the following criteria: p < 0.05. BME, basement membrane extract; BM MSC, bone marrow-derived mesenchymal stem cell; Cart-Org, cartilage-organoid.

Chondrocyte differentiation and hypertrophy are positively regulated by signaling molecules, including TGFβ/BMPs, Parathyroid hormone-related protein (PTHrP), Indian hedgehog (IHH), fibroblast growth factors (FGFs), and Wnt (12–14). TGFβ/BMPs induces SMAD pathway activation, which upregulates a chondrogenic master regulator, SOX9, in the early phase of chondrocyte differentiation. PTHrP, IHH, FGFs, and Wnt transduce chondrocyte maturation signals during pre-hypertrophy/hypertrophy. The IPA data indicated that the SMAD pathway, especially SMAD1-related signaling, was an activated upstream regulator (Fig. 4A and Fig. S2). SMAD1 forms a complex with SMAD5 and SMAD9 (SMAD1/5/9), which is phosphorylated via the activation of type I receptors ALK 3 (BMPR1A) or ALK 6 (BMPR1B) in conjunction with the type II receptor BMPR2. IPA also predicted that ALK3 (BMPR1A) is an activated upstream regulator in the BM MSC-based Cart-Org with BME. Given these observations, we consider that BME contains some type of BMPR1A/BMPR2 ligands, for example, BMPs, resulting in the activation of the BMPR1A/BMPR2-SMAD1/5/9 pathway.

The nuclear factor kappa-light-chain-enhancer of activated B cells (NF-κB) negatively regulates cartilage differentiation (15). Activation of the NF-κB pathway induced by inflammatory cytokines, such as IL-1 and TNF, suppresses the gene expression of SOX9 and cartilage-specific ECMs, including aggrecan, collagen type II, collagen type IX, and collagen XI (16–18). The present IPA data showed that IL1B and TNF were the inhibited upstream regulators. The members of the NF-κB family, NFKB1 (p105/50) and RELA (p65), were prominently observed in the pathways of inhibited upstream regulators (Fig. S3). These results indicate that BME further promotes cartilage development of the BM MSC-based Cart-Org by inhibition of the NF-κB pathway, whereas its particular substance(s) within the BME has not yet been identified.

IPA downstream effects analysis using a dataset of BM MSC-based Cart-Org on day 7 predicted that cartilage development was a significant biological downstream activity upon BME stimulation (Fig. 4B and C, significance criteria: p < 0.05, Z-score > 2). In addition, gene ontology (GO) analysis using a dataset of the BM MSC-based Cart-Org on day 21 further indicated that several gene sets of biological processes, skeletal system development, ossification, and cartilage development were significantly enriched in the BM MSC-based Cart-Org with BME on day 21 (Fig. 4D, significance criteria: p < 0.05). Transcripts associated with the extracellular region, extracellular space, and growth factor activity were also enriched in the BM MSC-based Cart-Org with BME on day 21. Briefly, comprehensive transcriptome analyses indicated that the BME-mixed 3D pellet culture facilitated progressive chondrogenesis and further ossification of the BM MSC-based Cart-Org. In addition, BME modulates several intracellular signaling during the early chondrogenic phase of the BM MSC-based Cart-Org, notably the pathways of SMAD and NF-κB.

### BME stimulation upregulates gene expression associated with endochondral ossification

Given that BME promotes cartilage development and ossification of the Cart-Org, we further examined the time transition of the expression of each gene associated with endochondral ossification (4,19,20). The representative chondrogenesis/ossification gene expression transitions are summarized in Fig. 5A. During early chondrogenesis, *SOX9*, *COL2*, and *ACAN* are expressed in chondrogenic progenitor/proliferating chondrocytes. During the phase transition from proliferating to pre-hypertrophic chondrocytes, fibroblast growth factor receptor 3 (*FGFR3*) and collagen type IX (*COL9*) are upregulated. Pre-hypertrophic chondrocytes were marked by the upregulation of *IHH* and Parathyroid hormone 1 receptor (*PTH1R*). At the latest chondrogenic stage, hypertrophic chondrocytes express collagen type X (*COL10*). Subsequently, hypertrophic chondrocytes convert their gene expression from cartilage anabolism to catabolism and ossification, which is caused by the upregulation of Sp7 Transcription Factor (*SP7*), alkaline phosphatase (*ALPL*), and matrix metalloprotease 13 (*MMP13*).

**Figure 5.**
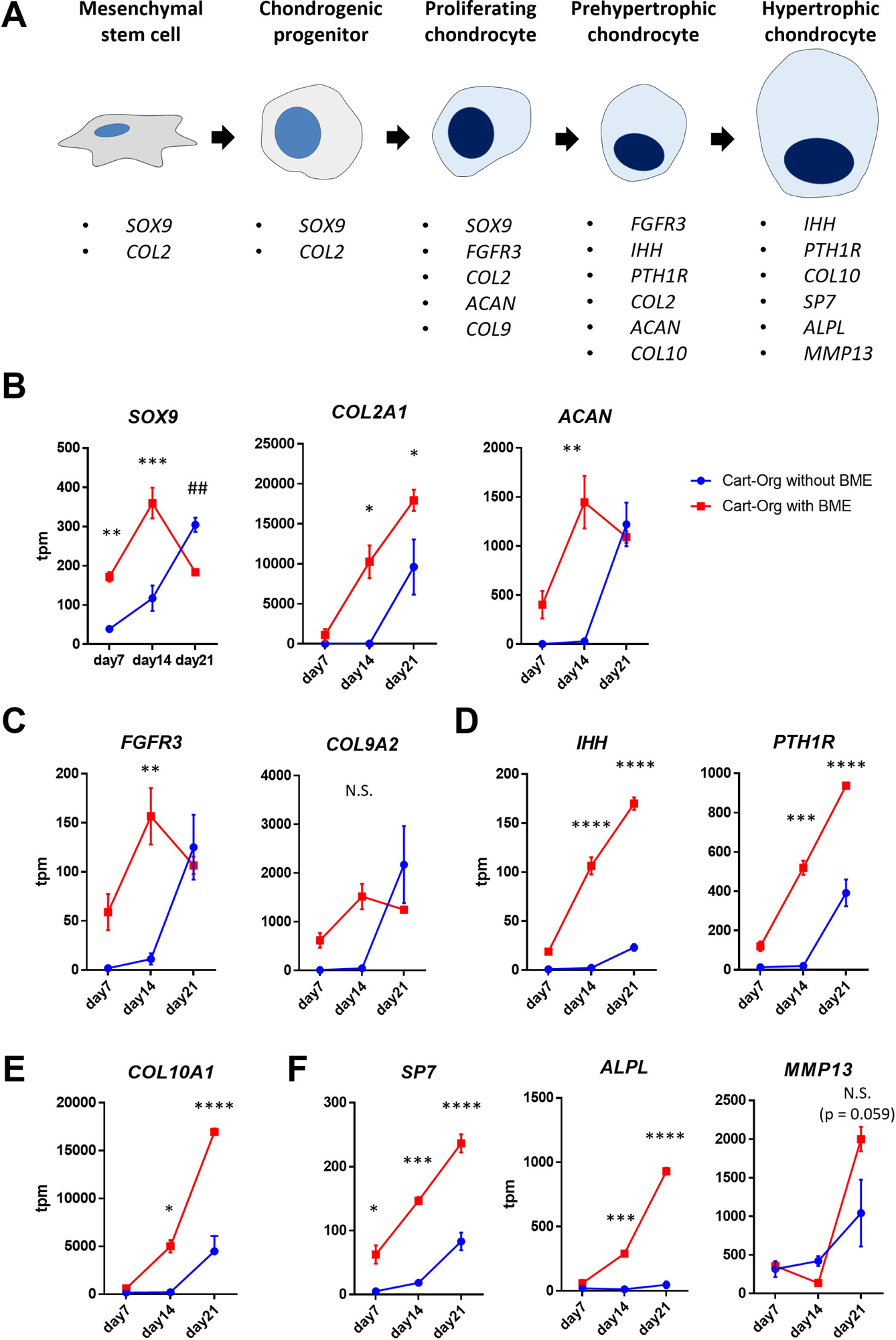
Time transition of endochondral ossification-associated gene expression in the BM MSC-based Cart-Org. A, the representative chondrogenesis/ossification gene-expression pattern in cartilage development. B–F, the transcript expression level of the gene associated with endochondral ossification. The transcript expression level of each gene is evaluated by transcripts per million (tpm) of the transcriptomic data in RNA-seq. The representative genes in the early chondrogenesis (B), the phase transition from the proliferating to the pre-hyperthophic chondrocyte (C), the pre-hypretrophic chondrocyte (D), the latest chondrogenic stage (E), and the phase of cartilage catabolism and ossification (F). *p < 0.05. **p < 0.01. ***p < 0.001. ****p < 0.0001. N.S. indicates a non-significant difference. Statistical analysis was performed by two-way ANOVA and Sidak’s multiple comparison test (n = 2 or 3 per group). The error bars represent SEMs. BM MSC, bone marrow-derived mesenchymal stem cell; SEM, standard error of the mean; Cart-Org, cartilage-organoid.

The genes associated with the early and transitional chondrogenic phases (*SOX9*, *COL2A1*, *ACAN*, *FGFR3*, *and COL9A1*) were highly expressed in the Cart-Org with BME on day 7 and were elevated on day 14, whereas the expression levels of these genes were low or undetectable in the Cart-Org without BME on days 7 and 14 (Fig. 5B and C). On day 21, the expression of these genes, except *COL2A1*, was downregulated in the Cart-Org with BME but upregulated in the Cart-Org without BME. *COL2A1* expression was progressively upregulated at a high level in Cart-Org treated with BME. The expression of genes associated with pre-hypertrophy (*IHH* and *PTH1R*) and hypertrophy (*COL10A1*) was markedly upregulated on day 14 and further elevated on day 21 in the Cart-Org with BME (Fig. 5D and E). Pre-hypertrophic/hypertrophic gene expression in the Cart-Org without BME was detected only on day 21. These data support the observation that BME promotes the chondrogenic phenotype of BM MSC-based Cart-Org.

In terms of gene expression related to cartilage catabolism and ossification, *SP7, ALPL,* and *MMP13* were highly expressed in the Cart-Org with BME when compared to those in Cart-Org without BME (Fig. 5F). SP7/Osterix is a key transcriptional determinant of osteoblast differentiation (21). Alkaline phosphatase is an enzyme involved in cartilage mineralization that removes extracellular pyrophosphate, an inhibitor of hydroxyapatite (22). MMP13, also known as collagenase-3, is a major enzyme that degrades cartilage by cleaving type II, type IX, and type X collagen and aggrecan (20,23). High expression and synthesis of MMP13 in hypertrophic chondrocytes induces the cartilage to enter a catabolic state, which results in marrow cavity formation during endochondral ossification. These data indicate that BME potentiated the endochondral ossification phenotype of the MSC-based Cart-Org.

## Discussion

Endochondral ossification is a developmental process in the skeletal system and bone marrow of vertebrates. Murine embryonic metatarsal explant cultures and rat femur slice cultures (organotypic culture) have been established as experimental models for endochondral ossification (24,25). An implant model was also developed, in which human cartilage derived from polydactyly epiphyseal chondrocytes was inoculated into the subcutaneous space of immunodeficient mice (26). However, these models depend on animal resources and polydactyly. To overcome these limitations, it is necessary to establish a stem cell-based Cart-Org with an endochondral ossification phenotype. In this study, we generated enlarged and mature human BM-MSC-based Cart-Orgs using a BME-mixed 3D pellet culture. BME modulated the gene expression pattern of the BM-MSC condensate to a chondrogenesis-dominant state, which promoted cartilage development and further ossification of the Cart-Org.

Endochondral ossification has been shown to occur via several cellular mechanisms (27). Canonical endochondral ossification is a mechanism by which hypertrophic chondrocytes undergo apoptosis and periosteum-derived osteoblasts populate the ossification center, followed by bone formation (19). Other mechanisms are based on chondrocyte transdifferentiation, such as chondrocyte-to-osteoblast transdifferentiation, dedifferentiation-to-redifferentiation, and direct differentiation. Chondrocyte-to-osteoblast transdifferentiation is a model in which immature chondrocytes differentiate into osteogenic precursors (28,29). This model appears to be specific to metaphysis formation in embryonic POC and postnatal bone growth. Dedifferentiation-to-redifferentiation is a model in which hypertrophic chondrocytes dedifferentiate into immature chondrocytes and subsequently redifferentiate into osteogenic lineages during POC and SOC development, as well as bone fracture healing (30–33). Direct differentiation is a cellular process in which hypertrophic chondrocytes directly differentiate into osteoblasts, which predominantly occurs during SOC development (34–36). It has been reported that SP7/Osx expression in the hypertrophic chondrocytes is a possible indicator of the direct differentiation (27). In the present study, the MSC-based Cart-Org with BME showed upregulation of *SP7* particularly with the late cultivation phase. This suggests that the Cart-Org with BME undergoes SOC-like endochondral ossification in association with direct differentiation.

The next challenge is the vascularization of the Cart-Org. Vascularized Cart-Org has the potential to be developed into a bone marrow regenerative model, that is, a bone marrow-organoid model. During endochondral ossification, vascular invasion is facilitated by the vasculogenic/angiogenic factor [vascular endothelial growth factor A (VEGFA)], which is secreted by hypertrophic chondrocytes (5,37,38). That is, Cart-Org must be highly expressed to secrete VEGFA for vascularization. We generated MSC-based Cart-Org with the endochondral ossification-dominant phenotype; however, BME-mixed 3D pellet culture did not substantially increase gene upregulation and secretion of VEGFA (Fig. S4). To augment VEGF production, Cart-Org must possess biological properties similar to those of the organism. Mechanical loading, such as hydrostatic shear or tensile stress, promotes cartilage ossification (39). Oxygen tension is also involved in cartilage development and ossification (12). These environmental and physical factors extensively contribute to the establishment of the bio-mimicking Cart-Org.

The stem cell-based endochondral ossification model would be beneficial not only for fundamental research in the field of skeletal system development but also for clinical applications, such as bone repair in fractures. For clinical application, Cart-Orgs must be produced under xeno-free culture conditions. In this study, we proposed that simultaneous regulation of the SMAD and NF-κB pathways caused Cart-Orgs to become enormous and mature. These findings suggest that cytokines and/or low-molecular-weight compounds that modulate these pathways may provide cues for the development of new xeno-free agents for Cart-Org generation.

In summary, we report that BME induces a predominant state in human BM-MSC condensate via activation of SMAD and inhibition of NF-κB, which leads to cartilage development and further ossification of the Cart-Org. The Cart-Org with the phenotype of endochondral ossification is supposed to possess potential for use in fundamental research on the skeletal system and marrow development, as well as in clinical applications. Further studies will provide more effective systems and agents for generating Cart-Org with an endochondral ossification phenotype.

## Experimental procedures

### Basement membrane extract (BME)

In this study, Matrigel Growth Factor Reduced Basement Membrane Matrix (Lot: 316001, 1257002, and 3032003, Corning) and Cultrex Reduced Growth Factor Basement Membrane Extract PathClear (Lot: 1675678, R&D) were used as the BME.

### Cell culture

Human bone marrow mesenchymal stem cells (BM MSCs) (TaKaRa Bio) were cultured in Mesenchymal Stem Cell Growth Medium 2 (TaKaRa Bio) supplemented with penicillin/streptomycin/amphotericin B (Wako). Accutase (Innovative Cell Technologies) was used as a cell-detachment agent for passaging. Cells were cryopreserved with Cryo-SFM (TaRaRa Bio), and long-term storage was performed in liquid nitrogen after slow freezing at -80°C.

### Chondrogenic differentiation in 3D pellet culture system

Human BM MSCs (5.0 × 10^5^ cells/15 mL centrifugation tube) were centrifuged at 300 × *g* for 5 min. After aspiration of the supernatant, the cells were resuspended with 0.5 mL of Complete MesenCultTM-ACF Chondrogenic Differentiation Medium (Stem Cell Technology) supplemented with antibiotics and centrifuged at 300 × *g* for 10 min to obtain a cell pellet. The tubes were placed vertically in an incubator at 37°C and 21% O_2_. Chondrogenic differentiation was performed for 21 days with a culture medium exchange every 3 days.

In the case of BME-mixed 3D pellet culture, human BM MSCs (5.0 × 10^5^ cells/15 mL centrifugation tube) were centrifuged at 300 × *g* for 5 min. After aspiration of the supernatant, the cell pellet was loosened by tapping and mixed with 10 μL BME. The cell/BME mixture was allowed to gel at 37°C for 15 min, and 0.5 mL of Complete MesenCultTM-ACF Chondrogenic Differentiation Medium was added. Chondrogenic differentiation was performed for 21 days with a culture medium exchange every 3 days.

### Preparation of frozen sections

BM-MSC-based artificial cartilages (cartilage organoids: Cart-Orgs) were washed with phosphate-buffered saline (PBS) and fixed in PBS-based 4% paraformaldehyde (4% PFA, Wako) for at least 24 h. The fixed samples were subsequently immersed in PBS-containing 30% sucrose (Wako, Osaka, Japan) for cryoprotection. Treated Cart-Orgs were embedded in OCT (Tissue-Tek, Sakura Finetek Japan) at -80°C. Frozen samples were sliced at 10 µm thin-section using a cryostat (Leica Biosystems).

### Hematoxylin and Eosin (HE) and Alcian blue staining

The frozen sections were post-fixed with a fixing solution (100% ethanol, formalin stock solution, and acetic acid) and washed with diluted water. For HE staining, the slides were sequentially immersed in hematoxylin solution (Muto Kagaku, Tokyo, Japan) for 1 min, warm water for hematoxylin staining, eosin solution (Muto Kagaku) for 5 s, and water for rinsing. For Alcian blue staining, the slides were first immersed in 3% acetic acid to equilibrate the pH of the sections. After soaking the slides in an Alcian blue staining solution (Muto Kagaku) for 1 min, they were washed with 3% acetic acid and rinsed with water. The slides were counterstained with Fast Red (ScyTek Laboratories, Logan, Utah, USA). After staining, the slides were passed several times through the baths in the order of 70% ethanol, 95% ethanol, and 100% ethanol for dehydration and immersed in 100% ethanol for 5 min. The slides were then passed through four Lemosol baths (Wako, Osaka, Japan). The slides were sealed in Multimount 480 (Matsunami Glass Industry, Osaka, Japan).

### RNA-seq

Total RNA isolation, RNA-seq library preparation, and RNA-seq were performed by Rhelixa (Tokyo, Japan). Total RNA from the Cart-Org was isolated using ISOSPIN Cell & Tissue RNA (NIPPON GENE). The RNA integrity score was calculated using NanoDrop One (Thermo Fisher Scientific) in TapeStation 4150 (Agilent Technologies). RNA-Seq libraries were prepared with NEBNext Poly(A) mRNA Magnetic Isolation Module and NEBNext® UltraTMII Directional RNA Library Prep Kit (New England Biolabs). Libraries were sequenced using the NovaSeq 6000 system (Illumina) with 150bp×2 paired-end reads.

### RNA-seq data analysis

RNA-seq data analysis was outsourced to Rhelixa. The quality of raw paired-end sequence reads was assessed using FastQC (version 0.11.7). Low-quality (< 20) bases and adapter sequences were trimmed using the Trimmomatic software (version 0.38) with the following parameters: ILLUMINACLIP: path/to/adapter.fa:2:30:10 LEADING:20 TRAILING:20 SLIDINGWINDOW:4:15 MINLEN:36. The trimmed reads were aligned to the reference genome (GRCh38/hg38) using the RNA-seq aligner HISAT2 (version 2.1.0). The HISAT2-resultant .sam files were converted to .bam files using Samtools (version 1.9), and the .bam files were used to estimate the abundance of uniquely mapped reads using FeatureCounts (version 1.6.3). Raw read counts were normalized to transcripts per million (TPM). Heatmaps and volcano plots were created from the Z-scores of the normalized counts using the Stats (version 3.6.1) and gplots (version 3.0.1.1) R packages. PCA of the normalized counts was conducted, and each sample was projected onto the 2D plane of the first and second PCA axes using the Stats (version 3.6.1) and gplots (version 3.0.1.1) R packages. Raw read counts were normalized by relative log normalization, and differentially expressed genes were analyzed using DESeq2 (version 1.24.0). Differentially expressed genes (DEGs) were detected with the thresholds of |log2FC (Fold Change)| > 1 and adjusted p value < 0.05, using the Benjamini and Hochberg (BH) method.

### Ingenuity pathway analysis (IPA)

IPA (Ingenuity Systems, QIAGEN) was performed to identify significant upstream regulators, signaling pathways, and significant biological downstream activities in the BM MSC-based Cart-Org with BME on day 7.

### Gene ontology (GO) enrichment analysis

GO enrichment analysis was performed using the Rhelixa database. The dataset of DEGs between the BM MSC-based Cart-Org with or without BME on day 21 was analyzed using GOATOOLS (version 1.1.6). The p-value was corrected using the Benjamini-Hochberg method for multiple testing calibrations.

### Protein array

Angiogenic regulators profiled in the conditioned medium were analyzed using a Proteome Profiler Human Angiogenesis Array Kit (ARY007; R&D Systems). The conditioned medium (1 mL) was loaded onto the array membrane, according to the manufacturer’s instructions. ECL Prime (GE Healthcare) was used as the horseradish peroxidase substrate, and chemiluminescence signals were detected using ChemiDoc (BIO RAD). For the quantitative spot index, mean pixel intensity (MPI) was calculated using ImageJ 1.46r.

### Statistical analysis

Quantitative data are depicted as mean ± standard error values of the mean (SEM). Two-group comparisons were performed using an unpaired Student’s t-test. Multigroup comparisons were performed using two-way analysis of variance (ANOVA) and Sidak’s multiple comparison test. Statistical analyses were performed using GraphPad Prism 6 (GraphPad Software).

## Data availability statement

The data generated in this study are available from the corresponding author upon reasonable request.

## Supporting information

This article contains supporting information.

## Acknowledgments

### Author contributions

H.N., S. Y., and S. T. methodology; H.N., S. Y., and S. T., investigation; H. N., S. Y., Nobuaki. S., A. S., S. O., and S. T. formal analysis; H. N and S.T. writing–original draft; Nobuaki S., A. S., S. O., T. Kanematsu., Naruko S. A. K., T. Kojima, T. M., and S. T. supervision; Nobuaki S., T. Kojima, T. M., and S. T. conceptualization; H. N., S. Y., Nobuaki S., A. S., S. O., T. Kanematsu., Naruko S. A. K., T. Kojima, T. M., and S. T. writing–review and editing; H. N., S. Y., Nobuaki S., A. S., S. O., T. Kanematsu., Naruko S. A. K., T. Kojima, T. M., and S. T. data curation; Nobuaki S., A. K., T. Kojima, T. M., and S. T. funding acquisition; S. T. validation; S. T. visualization; S. T. project administration.

### Funding

This study was supported by grants-in-aid provided by the Japanese Ministry of Education, Culture, Sports, Science, and Technology (Grant No. 17H05073 to S. T. and Grant No. 19K08853 to A. K.), the National Center for Geriatrics and Gerontology (NCGG, the Research Funding for Longevity Sciences, Grant No. 22-10 to A. K.), the Takeda Science Foundation (to S. T.), the SENSHIN Medical Research Foundation (to S. T.), and JST FOREST Program (Grant No. JPMJFR2158 to S. T.).

## Supporting information

**Figure S1. Volcano plot of the differentially expressed genes in RNA-seq on the BM MSC-based Cart-Org with BME.** A–C, the transcriptomic datasets were processed by volcano plot analysis to graphically compare the upregulated and downregulated genes of the BM MSC-based Cart-Org with BME against the BM MSC-based Cart-Org without BME. Each plot is displayed as day 7 (A), day 14 (B), and day 21 (C) of culture. The significance of the data was determined via the following criteria: p < 0.05, fold change > 2, or < -2. Blue, gray, and red plots indicate downregulated, non-differentiated, and upregulated genes, respectively. RNA-seq, RNA sequencing; BME, basement membrane extract; BM MSC, bone marrow-derived mesenchymal stem cell; Cart-Org, cartilage-organoid.

**Figure S2. The pathways of upstream regulators with activated state in the BM MSC-based Cart-Org with BME on day 7.** The pathways are shown as signaling toward *SOX9, COL2A1*, and *ACAN*. BME, basement membrane extract; BM MSC, bone marrow-derived mesenchymal stem cell; Cart-Org, cartilage-organoid.

**Figure S3. The pathways of upstream regulators with inhibited state in the BM MSC-based Cart-Org with BME on day 7.** The pathways are shown as signaling toward *SOX9, COL2A1*, and *ACAN*. BME, basement membrane extract; BM MSC, bone marrow-derived mesenchymal stem cell; Cart-Org, cartilage-organoid.

**Figure S4. BME-mixed 3D pellet culture did not increase the gene upregulation and secretion of VEGFA.** A, *VEGFA* expression level in the BM MSC-based Cart-Org with or without BME on each culture day. The transcript level is evaluated by transcripts per million (tpm) of the transcriptomic data in RNA-seq. N.S. indicates a non-significant difference. Statistical analysis was performed by two-way analysis of variance (ANOVA) and Sidak’s multiple comparison test (n = 2 or 3 per group). The error bars represent SEMs. B, the profile of angiogenic factors in the culture medium of the BM MSC-based Cart-Org. The angiogenic factors were profiled using the Proteome Profiler human Angiogenesis Array Kit. The top and bottom panels indicate the array images generated with the Cart-Org without or with BME, respectively. The spots of VEGFA are pointed out by red rectangles. VEGFA, vascular endothelial growth factor A; MPI, mean pixel intensity; RNA-Seq, RNA sequencing; BME, basement membrane extract; BM MSC, bone marrow-derived mesenchymal stem cell; Cart-Org, cartilage-organoid; SEM, standard error of the mean.

