## Supporting Information for "Basement membrane extract potentiates the endochondral ossification phenotype of bone marrow-derived mesenchymal stem cell-based cartilage organoids"

List of supporting information:

1. Supporting figures (Figures S1–S4)

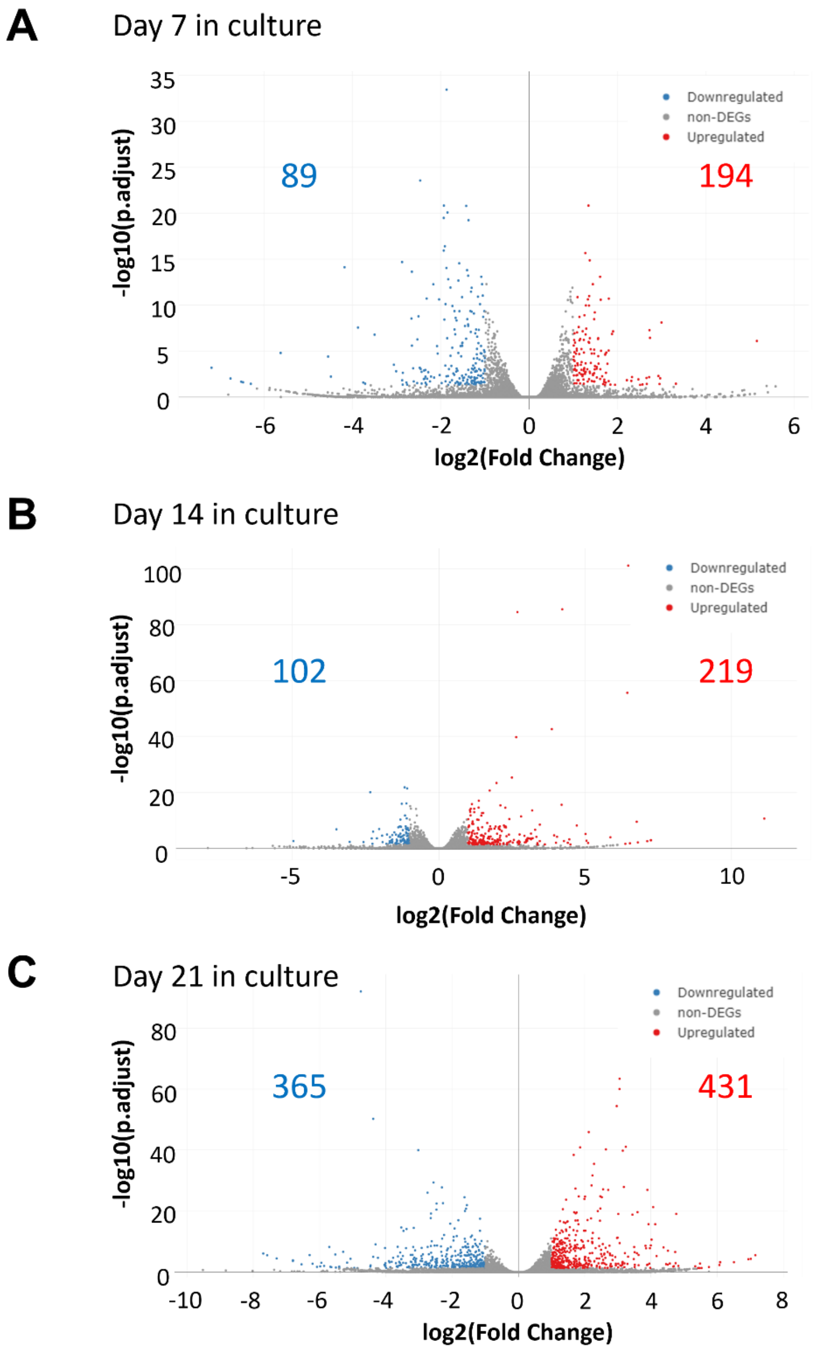


**Figure S1. Volcano plot of the differentially expressed genes in RNA-seq on the BM MSC-based Cart-Org with BME.** A–C, the transcriptomic datasets were processed by volcano plot analysis to graphically compare the upregulated and downregulated genes of the BM MSC-based Cart-Org with BME against the BM MSC-based Cart-Org without BME. Each plot is displayed as day 7 (A), day 14 (B), and day 21 (C) of culture. The significance of the data was determined via the following criteria: $p < 0.05$, fold change > 2, or < -2. Blue, gray, and red plots indicate downregulated, non-differentiated, and upregulated genes, respectively. RNA-seq, RNA sequencing; BME, basement membrane extract; BM MSC, bone marrow-derived mesenchymal stem cell; Cart-Org, cartilage-organoid.

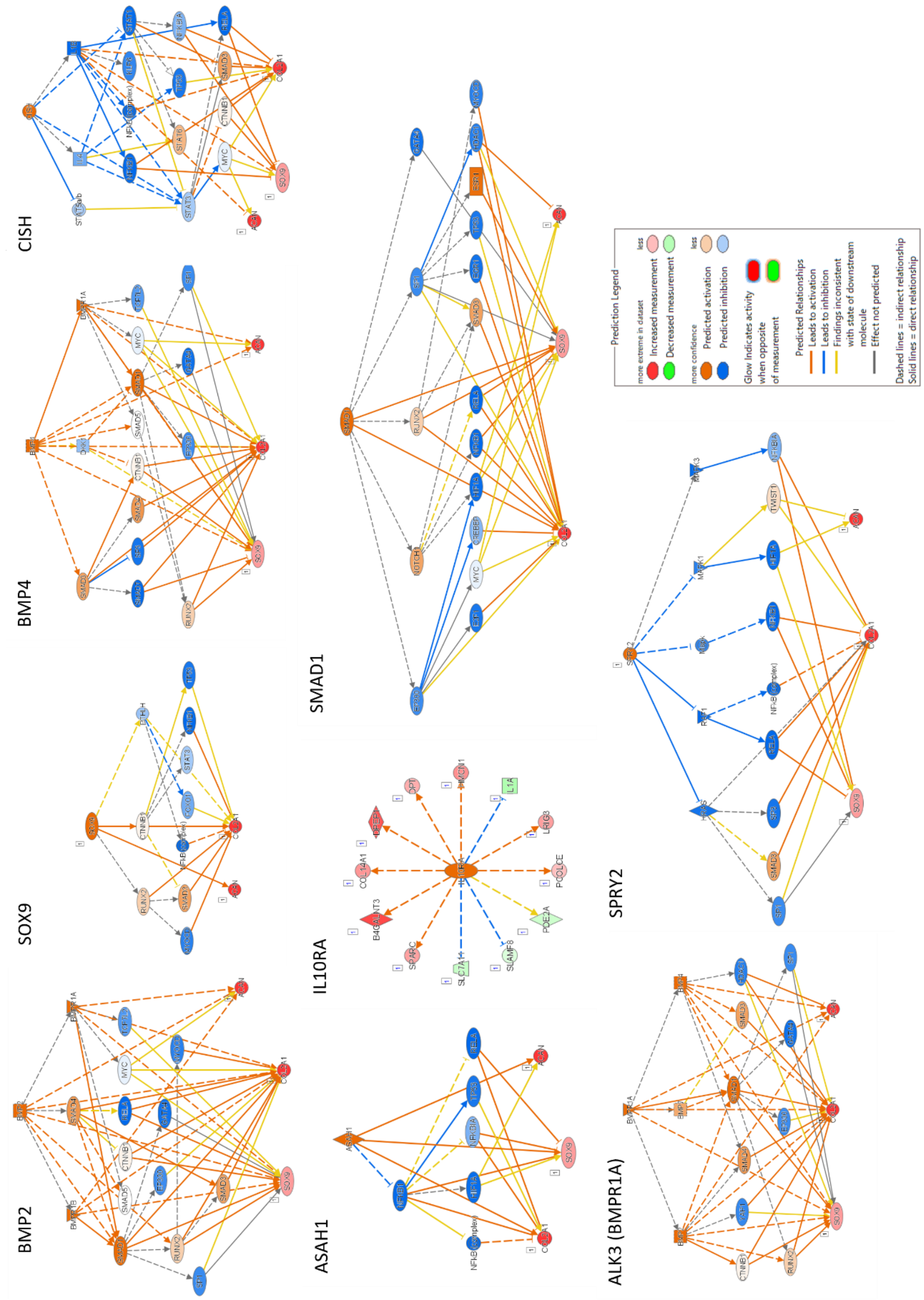


**Figure S2. The pathways of upstream regulators with activated state in the BM MSC-based Cart-Org with BME on day 7.** The pathways are shown as signaling toward *SOX9, COL2A1*, and *ACAN*. BME, basement membrane extract; BM MSC, bone marrow-derived mesenchymal stem cell; Cart-Org, cartilage-organoid.

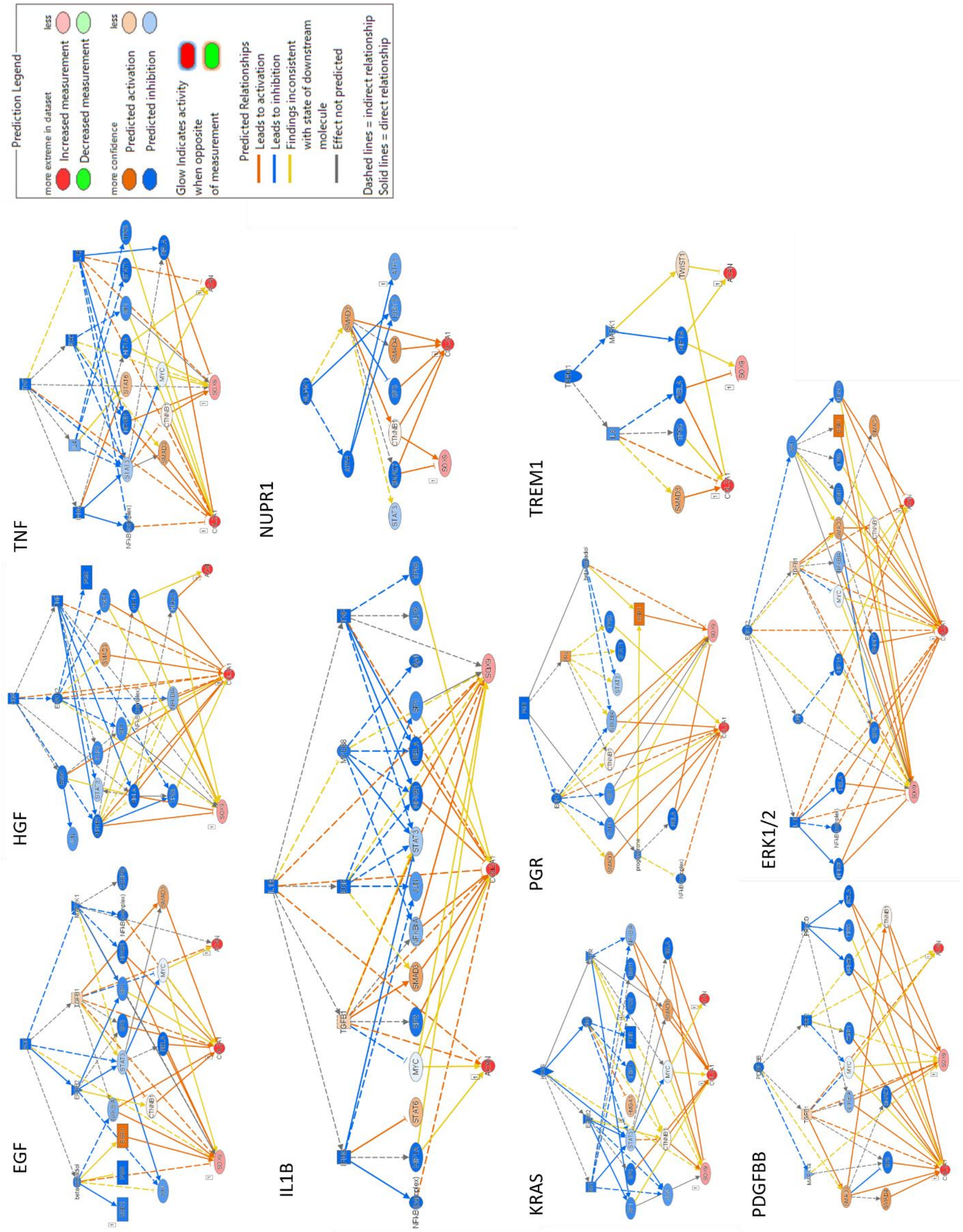


**Figure S3. The pathways of upstream regulators with inhibited state in the BM MSC-based Cart-Org with BME on day 7.** The pathways are shown as signaling toward *SOX9, COL2A1*, and *ACAN*. BME, basement membrane extract; BM MSC, bone marrow-derived mesenchymal stem cell; Cart-Org, cartilage-organoid.

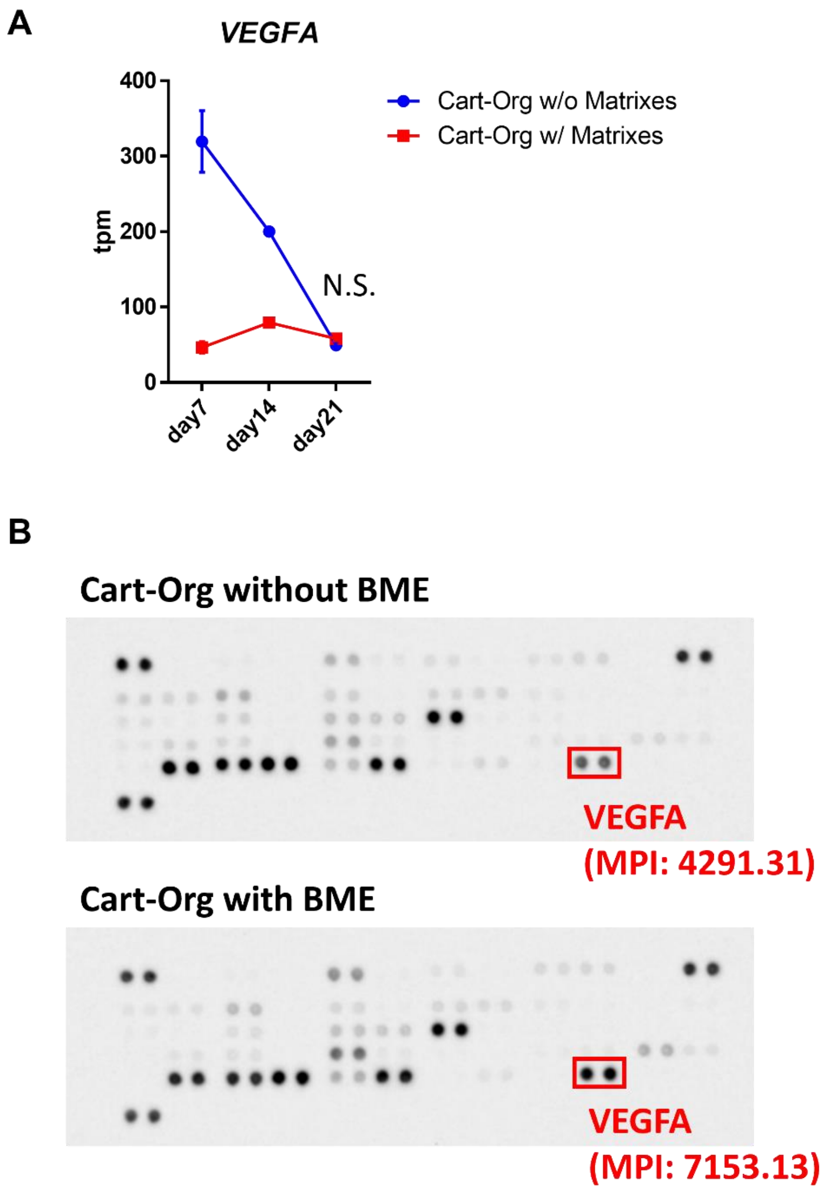


**Figure S4. BME-mixed 3D pellet culture did not increase the gene upregulation and secretion of VEGFA.** A, *VEGFA* expression level in the BM MSC-based Cart-Org with or without BME on each culture day. The transcript level is evaluated by transcripts per million (tpm) of the transcriptomic data in RNA-seq. N.S. indicates a non-significant difference. Statistical analysis was performed by two-way analysis of variance (ANOVA) and Sidak's multiple comparison test (n = 2 or 3 per group). The error bars represent SEMs. B, the profile of angiogenic factors in the culture medium of the BM MSC-based Cart-Org. The angiogenic factors were profiled using the Proteome Profiler human Angiogenesis Array Kit. The top and bottom panels indicate the array images generated with the Cart-Org without or with BME, respectively. The spots of VEGFA are pointed out by red rectangles. VEGFA, vascular endothelial growth factor A; MPI, mean pixel intensity; RNA-Seq, RNA sequencing; BME, basement membrane extract; BM MSC, bone marrow-derived mesenchymal stem cell; Cart-Org, cartilage-organoid; SEM, standard error of the mean.
